# Co-exposure to multiple *Ranavirus* types enhances viral infectivity and replication in a larval amphibian system

**DOI:** 10.1101/329821

**Authors:** Joseph R. Mihaljevic, Jason T. Hoverman, Pieter T.J. Johnson

## Abstract

Multiple pathogens commonly co-occur in animal populations, yet few studies demonstrate how co-exposure of individual hosts scales up to affect transmission. Although viruses in the genus *Ranavirus* are globally widespread and multiple virus species or strains likely co-occur in nature, no studies have examined how co-exposure affects infection dynamics in larval amphibians. We exposed individual *Rana aurora* (Northern red-legged frog) larvae to *Ambystoma tigrinum* virus (ATV), frog virus 3 (FV3), or an FV3-like strain isolated from a frog-culturing facility in Georgia, USA (RCV-Z2). We compared single-virus to pairwise co-exposures, while experimentally accounting for dosage. Co-exposure to ATV and FV3-like strains resulted in almost twice as many infected individuals compared to single-virus exposures, suggesting an effect of co-exposure on viral infectivity. The viral load in infected individuals exposed to ATV and FV3 was also higher than the single-dose FV3 treatment, suggesting an effect of co-exposure on viral replication. In a follow-up experiment, we examined how the co-occurrence of ATV and FV3 affected epizootics in mesocosm populations of larval *Pseudacris triseriata* (Western chorus frog). Although ATV did not generally establish within host populations (<4% prevalence), when ATV and FV3 were both present, this co-exposure resulted in a larger epizootic of FV3. Our results emphasize the importance of multi-pathogen interactions in epizootic dynamics and have management implications for natural and commercial amphibian populations.

## Introduction

Classic theory and empirical research on infectious disease, in both wildlife and humans, has predominantly focused on the interaction between a single host and a single pathogen (Anderson et al. 1992, Hudson et al. 2002, Keeling & Rohani 2008, Rigaud et al. 2010, Tompkins et al. 2010). While substantial biological insights have been derived from such studies, multiple pathogens often co-occur (Petney & Andrews 1998, Pedersen & Fenton 2006, Balmer & Tanner 2011, Knowles et al. 2013, Griffiths et al. 2014, Stutz et al. 2018) and can result in transmission dynamics that deviate from classical expectations (Alizon et al. 2013a, Johnson et al. 2015, Seabloom et al. 2015), especially if host individuals become simultaneously or sequentially infected with different pathogens (i.e. co-infected or super-infected, respectively).

Research in a variety of systems has shown that ecological interactions among pathogens within a host, such as priority effects, competition, and facilitation, alter pathogen replication rates, probability of infection, clearance rates, and host survival (de Roode et al. 2005, Pedersen & Fenton 2006, Johnson & Hoverman 2012, Johnson, Preston, Hoverman, & LaFonte 2013, Nunn et al. 2014, Seabloom et al. 2015). While modeling studies have demonstrated how these within-host dynamics can scale up to affect transmission dynamics within host populations (Mideo et al. 2008, Alizon 2013), empirical studies linking scales in natural systems are limited. A notable exception involves de-worming experiments in wild buffalo populations, which show that co-infection with nematodes and the bacterial agent of tuberculosis increases host mortality (Jolles et al. 2008, Ezenwa & Jolles 2015). In an epidemiological model of the system, increased removal of hosts due to co-infection limited tuberculosis transmission in a manner consistent with large-scale epidemiological patterns in the field (Jolles et al. 2008, Ezenwa & Jolles 2015). Understanding how pathogen co-exposure affects pathology and transmission requires more studies that explore the impacts of pathogen co-exposure across multiple biological scales (Mihaljevic 2012, Gog et al. 2014, Buhnerkempe et al. 2015, Johnson et al. 2015).

Viruses of the genus *Ranavirus* (family: Iridoviridae) provide a tractable and relevant model system for exploring the effects of multiple pathogens at both the within-and among-host spatial scales. Ranaviruses infect amphibian communities globally and can cause massive die-off events (up to 100% mortality), constituting a major threat to wild and commercially maintained amphibian populations (Gray et al. 2009b, Lesbarrères et al. 2012, Gray & Chinchar 2015). There are several reasons to suspect that co-exposure to multiple *Ranavirus* types (e.g. viral species or strains) could be common in nature and influence transmission dynamics. First, this viral genus is genetically and ecologically diverse, with different type species and strains that show variability in epidemiological traits. For instance, two species of the genus common to North America – *Ambystoma tigrinum virus* (ATV) and *Frog virus 3* (FV3) – can easily be differentiated based on genomic characteristics, but also by their variability in infectivity, with ATV being more host-specific to salamanders (urodeles) and FV3 being more host-generalist, capable of infecting amphibians, reptiles, and some fish (Chinchar et al. 2009, 2011, 2017). Furthermore, unique strains of ATV and FV3 differ in the rates at which they cause host mortality, which we refer to as virulence (Brunner & Collins 2009, Hoverman et al. 2010). Finally, both ATV and FV3 can be highly prevalent across the landscape, and their spatial distributions broadly overlap (Tornabene et al., Gray et al. 2007, Ridenhour & Storfer 2008, Greer et al. 2009, Brunner et al. 2011, Hoverman, Mihaljevic, et al. 2012, Gray & Chinchar 2015), suggesting a high potential for co-occurrence. To date, however, no studies have considered the effect of multiple *Ranavirus* species or strains on disease outcomes or epizootics.

While ATV has high infectivity in salamanders (Picco et al. 2007, Brunner & Collins 2009), there is mixed evidence that strains of ATV are able to infect anuran (frog and toad) larvae (Jancovich et al. 2001, Schock et al. 2008). For example, of the three frog species experimentally exposed to ATV by Schock et al. (2008), all three species showed susceptibility to ATV infection, and a small proportion of individuals died of ATV-induced disease. However, Jancovich et al. (2001) exposed two frog species to ATV – including a different population of one species also studied by Schock et al. (2008) – and found no signs of infection. Together, these data suggest that ATV infections in anurans are possible, though the probability of infection likely varies among species and populations, and possibly among ATV strains. FV3, in contrast, shows high infectivity among diverse host species, and infection often leads to mortality in both anuran and salamander larvae (Brunner et al. 2005, Picco et al. 2007, Schock et al. 2008, Hoverman et al. 2010, 2011). Given that anurans and salamanders often co-occur as larvae (Hoverman, Gray, et al. 2012, Johnson, Preston, Hoverman, & Richgels 2013) and that ranaviruses infect multiple host species, it is likely that co-exposure and subsequent within-host interactions between virus types is relatively common in nature.

Here, we examined the effects of co-exposure to ATV and FV3 on mortality and transmission dynamics in larval amphibians. We conducted two experiments to assess how the effects of co-exposure scale-up from within-host outcomes to between-host transmission, ultimately affecting epizootics. Theory suggests that the effect on co-exposure on transmission will depend on disease outcomes and pathogen replication within hosts (Pedersen & Fenton 2006, Jolles et al. 2008, Mideo et al. 2008, Ezenwa & Jolles 2011). For instance, if within-host interactions lead to more rapid host death, then transmission of co-occurring pathogens could be dampened, leading to smaller epizootics. However, if co-exposure facilitates the invasion of pathogens, enhances within-host replication, or increases host tolerance to infection, transmission could be enhanced, leading to larger epizootics (Mideo et al. 2008, Ezenwa & Jolles 2011, Sofonea et al. 2015). *A priori* we expected that, due to the generally high infectivity and virulence of both ATV and FV3, co-exposure would increase host mortality and therefore limit epizootic size. To test these expectations, we performed experimental infections at two scales. We first exposed larval frogs individually to one or two virus types to determine how co-exposure affected mortality rate, the probability of infection, and within-host viral replication. We then conducted an experiment using replicate populations of larval frogs. Here, we pre-exposed larvae to either FV3 or ATV, and we added these individuals to small populations of susceptible larval frogs to explore how co-exposure affected the proportion of individuals infected and the average viral load. Our results indicated that co-exposure enhances viral infectivity and viral replication, illustrating the need to further explore how *Ranavirus* types are distributed across the landscape and how this might affect epizootics.

## Materials and Methods

### Viruses and culturing

Aliquots of *Ambystoma tigrinum* virus (ATV; Regina ranavirus (RRV) #11800) and frog virus 3 (FV3; #061405) were generously provided by V. Gregory Chinchar. The RRV strain of ATV was originally isolated in 1997 from *Ambystoma tigrinum* in Regina, Saskatchewan, Canada (Bollinger et al. 1999), and the FV3 strain is also a wild-type strain isolated from *Rana pipiens* populations of the Midwestern United States in the 1960’s (Granoff et al. 1965). An aliquot of the *Rana catesbeiana* virus (RCV-Z2) strain of FV3 (hereafter referred to as R-FV3) isolated from a ranaculture facility in Georgia in 2006, was generously provided by Matthew Gray and Debra Miller (GenBank accession no. EF101698; Miller and Rajeev 2007, Claytor et al. 2017). This strain was the cause of a die-off event in the facility’s bullfrog population, and is twice as virulent as wild-type FV3 in some amphibian species (Hoverman et al. 2010). We propagated the three viruses through immortalized fathead minnow (FHM) cells fed with Eagle’s minimum essential medium (MEM) with Hank’s salts, containing 5% fetal calf serum. Titer of the resulting viral stocks was determined by plaque assays using serial dilutions of the stock, resulting in titers represented in plaque forming units (PFU). It is important to note that we were unable to obtain an accurate titer of the R-FV3 stock before the start of the first experiment, which likely explains the observed lower-than-expected infectivity.

### Experiment 1: Individual-level

This experiment assessed the individual, host-level effects of co-exposure to ATV and FV3. Egg masses of *Rana aurora* were field-collected from wetlands in Oregon in spring 2012 and shipped to the University of Colorado at Boulder. Egg masses were first washed with sterile deionized water to remove any possible residual virions and were then placed into plastic containers for rearing. Larvae were reared at 20°C with a 12:12 hour day:night photoperiod and fed ground TetraMin^®^ fish flakes (Tetra) *ad libitum* until reaching Gosner stage 30 (Gosner 1960). At this time, larvae were randomly placed into individual, covered plastic containers (with drilled air holes) filled with 1-L of carbon-filtered, UV-sterilized water and allowed to acclimate for 24h. A subset of 15 larvae were euthanized by immersion in 1% buffered MS-222 and tested for infection to verify that none of the larvae harbored latent infections prior to experimentation (see quantitative PCR methods below). None of these individuals tested positive for ranaviruses.

Twenty-five larvae were assigned to each of 10 experimental treatment groups: a no-virus control, single dose of each virus alone (n=3 treatments), double dose of each virus alone (n=3 treatments), and each pairwise combination of the three viruses (i.e. a single dose of each of two viruses; n=3 treatments). Using this experimental design, we were able to account for additive and substitutive effects (e.g. dosage effects vs. effects of multiple strains). The control treatment consisted of a sham exposure to a 60μL aliquot of virus-free MEM. On 22 May 2012, a single dose (∼1×10^6^ PFU) or double dose (∼2×10^6^ PFU) of the respective virus or viruses was added to each larva’s container via sterile pipette tip. Thus, larvae were passively exposed to each virus inoculate, which likely better mimics natural transmission conditions relative to injection-based methods.

After virus addition, individuals were fed *ad libitum* every other day for the extent of the experiment. Complete water changes were conducted with carbon-filtered, UV-sterilized water every 4 days post-exposure (dpe) to ensure adequate water quality for the larvae. Standard protocols to avoid cross contamination between containers involved sterilizing dip nets with a 10% bleach solution for 10 minutes, followed by rinsing with sterile water to remove any residual bleach. Container and experimental room surfaces were cleaned with a 2% solution of Nolvasan between each container’s water changes, allowed to sit for 10 minutes, and then rinsed with sterile water.

The experiment ran for 21 d and mortality of larvae was monitored daily. If an individual died, the individual was extracted from its container, rinsed thoroughly with de-ionized water to remove any non-infecting virions that may have adhered to the individual’s skin, and then the entire individual was placed into a microcentrifuge tube and stored at −20°C for later processing. After 21d, all surviving larvae were euthanized in 1% buffered MS-222. These individuals were then washed thoroughly with de-ionized water, placed into individual microcentrifuge tubes, and stored at −20°C for later processing.

### Experiment 2: Population-level

A follow-up experiment tested how co-occurrence of ATV and FV3 in a larval amphibian population would affect transmission dynamics. Because the first, individual-level experiment showed qualitatively similar effects of co-exposure in the ATV+FV3 and ATV+R-FV3 treatments (see Results below), only ATV and FV3 were used for this experiment. In the spring of 2013, we were unable to obtain more *R. aurora* egg masses; instead, we collected egg masses of *Pseudacris triseriata* from local sites in Colorado, washed them with sterile deionized water, and reared them in plastic containers at 20°C with a 12:12 hour day:night photoperiod. Hatching larvae were fed ground TetraMin^®^ fish flakes (Tetra) *ad libitum* until reaching Gosner stage 30 (Gosner 1960).

The overall design of the experiment was to establish replicate populations of 10 uninfected larvae and then introduce 2 previously virus-exposed larvae into each population to track the spread of virus and determine if co-occurrence of ATV and FV3 alters the rate of spread and overall epizootic size. We used this method of transmission, instead of using passive exposure to MEM-suspended virus, because we wanted to assure that the behavior of infected hosts was allowed to affect transmission. To generate infected hosts for addition to the experimental populations, we randomly assigned a subset of the larvae to one of three exposure groups: FV3-exposure, ATV-exposure, and sham-exposure. Larvae were housed in 50-L covered plastic tubs (with drilled air holes) at densities no greater than 1 larva per liter of water. On 20 June 2013, larvae were passively batch-exposed to a dosage of 5×10^6^ PFU L^−1^ water of the respective virus or a sham exposure with an equivalent volume of virus-free MEM, using batches of 50 larvae. Larvae were held in these containers for 4d to initiate infection. In order to later identify which individuals were previously exposed, before adding the exposed individuals to the susceptible populations, we sedated each exposed individual and used a pair of micro-scissors to create a notch on the posterior, dorsal end of the tail. Unfortunately, these notches had healed by the end of the experiment and it was exceedingly difficult to identify which individuals were previously exposed.

Uninfected (i.e. susceptible) experimental populations were also established on 20 June 2013. We randomly selected 10 unexposed larvae and placed them in 15-L covered plastic tubs (with drilled air holes) filled with 12-L of carbon-filtered, UV-sterilized water. After the 4d batch-exposure, on 24 June 2013, each uninfected population received one of the following combinations of exposed larvae: (1) two sham-exposed larvae, (2) two FV3-exposed larvae, (3) two ATV-exposed larvae, or (4) one FV3-exposed larvae and one ATV-exposed larvae. Thus, each microcosm population contained 12 total *P. triseriata* larvae (10 susceptible and 2 exposed) for a total density 1 larva per liter of water. Each of the four treatments was replicated 6 times for a total of 24 experimental units.

We destructively sampled three replicates four days after the addition of the two exposed tadpoles (4dpe). This sample allowed us to establish an early epizootic time-point for comparison to late-stage epizootics. Larvae were extracted from each tub and individually euthanized in 1% buffered MS-222. As above, larvae were rinsed, placed into individual microcentrifuge tubes, and stored at −20°C for later processing. Starting at 5dpe, 80% water changes were implemented every 4 days for each remaining replicate. Mortality was continually monitored, and any deceased individuals were extracted from tubs, rinsed, and stored, as above. At 21dpe, the experiment was terminated and individuals processed as described above.

### Tissue processing and DNA extraction

Frozen samples were allowed to thaw to room temperature and 500μL of MEM was added to each microcentrifuge tube. The samples were then manually homogenized using a motorized homogenizer. This tissue homogenate was then centrifuged at 3000g for 1min. A 500μL aliquot of the resulting supernatant solution was placed into a new sterile microcentrifuge tube and used for DNA extraction. Qiagen™ DNeasy Blood and Tissue extraction kits and standard protocols were used to extract 250μL of buffered DNA suspension from each supernatant aliquot. DNA samples were stored at −20°C for later processing.

### Quantitative PCR amplification of viral DNA

The viral load of each DNA extract (in viral copy number equivalents) was evaluated using quantitative polymerase chain reaction (qPCR), estimated by comparison to a dilution series of standard DNA. We created a synthetic double-stranded DNA standard by synthesizing a 250bp fragment of the major capsid protein (MCP) gene (gBlocks^®^ Gene Fragments; Integrated DNA Technologies™), which is conserved among *Ranavirus* species (e.g. ∼97% sequence similarity between ATV and FV3 strains). We used a 10-fold dilution series from 2×10^8^ gene copies down to 2×10^1^ gene copies of standard DNA. Standards and samples were run in duplicate.

The qPCR protocol amplifies a ∼70bp region of the MCP, allowing the protocol to identify many *Ranavirus* species. However, importantly, the protocol cannot distinguish between virus species within a sample (Forson & Storfer 2006, Picco et al. 2007). Thus, we were unable to assess the simultaneous presence of virus types (i.e. co-infection). To test each sample for ranavirus infection, a 2.5μL volume of sample DNA was added to a reaction volume of 17.5μL containing the following reagents: 10μL TaqMan^®^ 2X Universal PCR Master Mix (No AmpErase UNG), 0.06μL forward primer (for a final concentration of 0.1μM; 5’ ACA CCA CCG CCC AAA AGT AC 3’), 0.18μL reverse primer (for a final concentration of 0.1μM; 5’ CCG TTC ATG ATG CGG ATA ATG 3’), 0.05μL fluorescent TaqMan^®^ probe (with a starting concentration of 100pmol μL^−1^; 5’ FAM-CCT CAT CGT TCT GGC CAT CAA CCA C-TAM 3’), and 7.21μL molecular grade water (Forson & Storfer 2006, Picco et al. 2007). All custom primers and probes were ordered through Life Technologies™. Samples were run in 96-well plates on an Applied Biosciences^®^ machine for 40 cycles: 95°C denaturing (20s), 54°C annealing (20s), and 72°C extension (30s). Two positive ATV and FV3 controls and two negative controls were run on each plate.

After qPCR analysis, the starting sample DNA concentrations of all virus-positive samples were estimated using a Quant-iT™ PicoGreen^®^ dsDNA Assay Kit (Life Technologies™). All viral loads were standardized to viral copy number per ng of sample DNA. We also quantified the DNA concentration of a random subset of non-infected samples in order to verify that viral detection was not dependent on a high concentration of initial sample DNA.

### Viral DNA sequencing of infected samples

We attempted to sequence a small region of the viral genome from all infected samples in order to verify the identity of the infecting virus(es). However, this method is not reliable at determining if multiple virus types are coinfecting an individual, especially if there are rare variants. Therefore, this analysis determined only the identity of the virus with the most DNA present in the sample (i.e. the most abundant virus in a given individual). Still, this sequencing helped narrow down the mechanisms driving the effects of co-exposure. We amplified a ∼350bp fragment of the MCP gene using a hemi-nested PCR protocol (Kattenbelt et al. 2000). The amplicon from each infected sample, along with a custom sequencing primer (5’ ACT ATG CCA CCT CCA TC 3’), was sent to Quintara Biosciences™ for Sanger sequencing. We also amplified and sequenced the same MCP gene fragment from the three *Ranavirus* strains used in the study (ATV, FV3, and R-FV3). We compared the sequencing data from each infected sample to that of the original viral strains.

### Statistical analyses: Experiment 1

All statistical analyses were conducted in the open-source software, *R* (R Core Development Team 2013). From the first experiment, we had three types of data for each of the 10 viral exposure treatments: survival, proportion of individuals becoming infected, and viral load per infected larva. We compared survival rates with a Mantel-Haenszel test, using the ‘*survival’* package. We first compared among single- and double-dose single-virus treatments, with dosage as the predictor variable. Because we found no difference in the survival rate between single- and double-dosages for the single-virus exposures 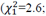 P = 0.11), we then compared the double-dose, single-virus treatments to the co-exposure treatments and control. A Cox proportional hazards model yielded the same qualitative results.

We conducted our analyses of proportion infected and viral load using the Bayesian statistical programming language, *Stan*, interfacing through *R* via the package ‘*rstan*’. We use this method because of the language’s flexibility in specifying the model structures for our analyses. In all cases, we used broad, vague priors for model parameters. We have made our code publicly available at the following link: https://bitbucket.org/jrmihalj/ranavirus_coexposure. To quantify the effects of virus identity, dosage, and co-exposure on the proportion of individuals that became infected, we conducted a logistic regression. Note that we conducted a similar analysis in the ‘*brglm*’ package in *R*, which uses a frequentist approach, and the results were qualitatively the same (not shown). The model that was fit in *Stan*, however, showed a more precise match between data and model predictions.

We structured our model as follows (using *R* linear model syntax for ease of interpretation): Number_Infected ∼ 0 + ATV + FV3 + RFV3 + Double: ATV + Double:FV3 + Double:RFV3 + FV3:RFV3 + ATV:RFV3 + ATV:FV3, where Number_Infected follows a binomial distribution, with *k* = 25 (i.e. the number of individuals exposed in each treatment). Thus, our model estimated a baseline effect of each virus type (i.e. a single-dose effect), a virus-specific effect of double dosage, and then the interactions between each virus type. These interactions represent the effect of co-exposure, which is an effect above and beyond the effect of a double dosage. Each one of model effects is compared to an intercept of zero, which on the logit scale equals 50% prevalence. Thus, for instance, a strong negative effect of ATV would mean that far less than 50% of individuals became infected with a single dose of ATV. In general, we were interested in whether the co-exposure effects are larger than all of the double-dose effects, indicating significant synergy between the two co-inoculating viruses.

Finally, to compare viral loads among treatments, we followed a similar approach as our treatment of the data on proportion infected. This model only used viral load data from the individuals that became infected. We therefore constructed a linear model predicting the natural log-transformed average viral copy number per ng DNA for each infected individual (averaged over the duplicate qPCR runs). We used similar model structure as above, except in this case we estimated an intercept representing the average infection intensity, and we did not include the single or double-dose ATV treatments, due to lack of any infections. We also added a term accounting for whether or not the individual died during the experiment. The model structure was therefore: Viral_Load ∼ Intercept + FV3 + RFV3 + Double:FV3 + Double:RFV3 + FV3:RFV3 + ATV:RFV3 + ATV:FV3 + Died. In this case, then, we were interested to see if the co-exposure groups had higher than average infection intensities, which again would indicate a synergistic effect of co-exposure.

### Statistical Analyses: Experiment 2

Similar to the first experiment, the second experiment had three response variables: survival rate, infection prevalence, and average viral load.However, this time, the response variables were population-specific, with 3 replicate populations per time-point (4dpe and 21dpe) and per treatment (FV3, ATV, FV3+ATV, Control). To compare survival among treatments, we used a Cox proportional hazards model with replicate population as a frailty term (i.e. analogous to random intercept term). We compared infection prevalence between the FV3 and FV3+ATV treatments by creating a generalized linear mixed effects model in *Stan* with prevalence explained by treatment, time (early vs. late), and their interaction. We excluded the ATV-only treatments due to the small number of infections and to simplify the analysis. The model was therefore of the form: Number_Infected ∼ Treatment + Time + Treatment*Time, where Number_Infected follows a binomial distribution with *k* = 12, the number of individuals in each replicate.

We similarly compared the viral load between the FV3 and FV3+ATV treatments by creating a linear mixed effects model with viral load (transformed as in the first experiment) explained by a treatment-by-time interaction, a fixed effect for whether the individual died or not, a fixed effect for day of death, and a random effect for replicate population. This model was of the form: Viral_Load ∼ Intercept + Treatment + Time + Treatment*Time + Died + Random(Replicate).

## Results

### Experiment 1

In the first experiment with *Rana aurora*, larvae experienced mortality throughout all treatments, including controls (Fig. A1). However, survival did not differ among treatments (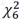 = 6.2; p = 0.40; Fig. A1). Infection prevalence in *R. aurora* ranged from 0-76%, with no individuals becoming infected in the sham control and no individuals becoming infected in the ATV-only treatments (including single- and double-doses of ATV).

Importantly, co-exposure to ATV and either FV3 or RFV3 caused a synergistic effect, enhancing overall infectivity compared to the double-dose treatments of single virus types (i.e. ATV, FV3, and RFV3 alone; Fig. 1a). Thus, the infection prevalence in co-exposure groups was nearly twice as high as single-virus exposures. However, co-exposure to both FV3-like strains (i.e. FV3+RFV3) did not cause such an effect (Fig. 1a). Indeed in the statistical model, the co-exposure effects of ATV:FV3 and ATV:RFV3 were larger than all other effects (Table 1), demonstrating a synergistic effect of co-exposure on viral infectivity.

**Table 1.** Effects of co-exposure on the proportion of *Rana aurora* hosts that became infected. The median and 95% credible interval (CI; the Bayesian analog to the 95% confidence interval) are shown for the coefficients in the logistic model. In general, effects whose 95% CI do not overlap zero are considered biologically meaningful. Co-exposure effects whose 95% CI do not overlap zero demonstrate an effect of co-exposure over and above any effects of dosage. These meaningful co-exposure effects are bolded.

| Effect |  | Median Coefficient | 95% CI |
| --- | --- | --- | --- |
| Baseline effects of each virus | ATV | -8.72 | (-20.03, -3.28) |
|  | FV3 | -1.44 | (-2.59, -0.50) |
|  | RFV3 | -0.78 | (-1.68, 0.08) |
| Effects of double-dose, compared to baseline | Double:ATV | 1.60 | (-7.53, 12.80) |
|  | Double:FV3 | 1.22 | (-0.01, 2.61) |
|  | Double:RFV3 | -0.17 | (-1.47, 1.03) |
| Co-exposure effects, above and beyond effects of dosage | FV3:RFV3 | 0.40 | (-1.04, 1.84) |
|  | **ATV:RFV3** | **8.52** | **(4.16, 16.75)** |
|  | **ATV:FV3** | **8.64** | **(4.32, 16.94)** |

**Figure 1.**
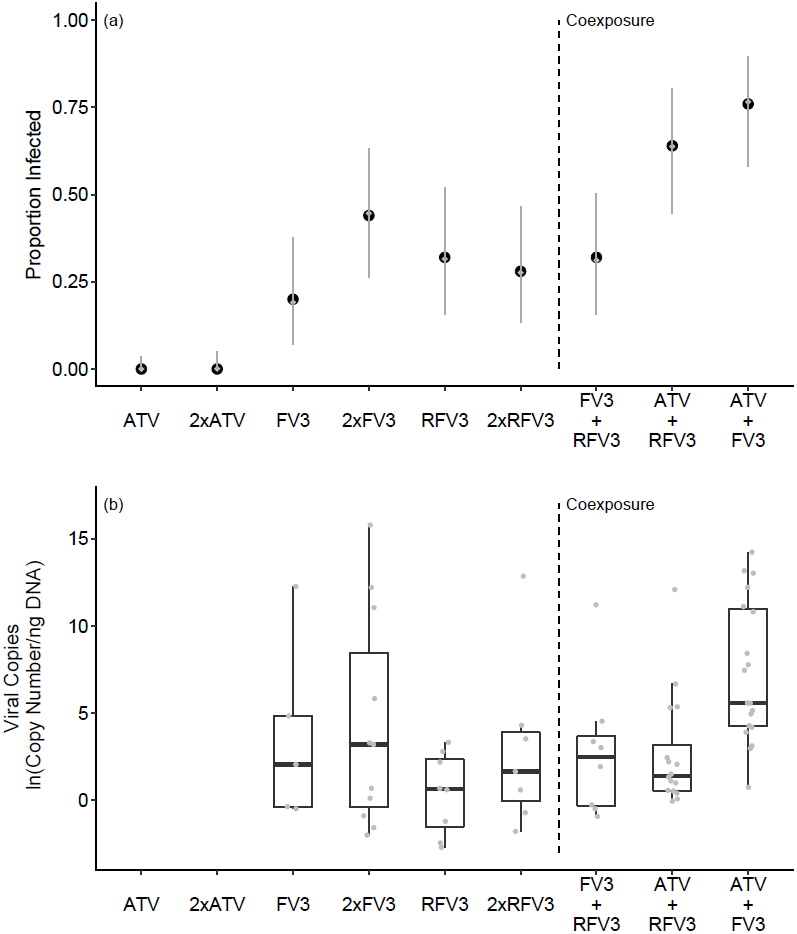
(a) Proportion of *Rana auora* individuals infected in all experimental groups from experiment 1. Gray points are the median predictions from the statistical model, and gray bars represent the estimated 95% credible intervals to show error in our estimates. (b) Boxplots of viral copy number per treatment group, with gray points representing the viral load of each measured host. The box represents the inter-quartile range (IQR; between first and third quartiles), and the center line marks the median value. The whiskers extend from the box to the highest or lowest value that is within 1.5 x IQR.

Infected individuals that died during the experiment had, on average, higher viral loads compared to infected individuals that survived to the end of the experiment (Table 2; Fig. A2). We also found that for RFV3, a double dose exposure led to a detectably higher average viral load compared to the single dose exposure, and there was a similar trend for FV3 (Fig 1b; Table 2). The viral loads of the ATV+FV3 co-exposed individuals were higher than the loads of individuals exposed to a single dose of FV3, but there was no difference in viral load in the ATV+FV3 treatment compared to exposure to a double dose of FV3 (Fig. 1b; Table 2). The other co-exposure treatments (ATV+RFV3 and FV3+RFV3) had similar viral loads to the single-dose exposure treatments of FV3 and RFV3 (Fig. 1b; Table 2).

**Table 2.** Effects of co-exposure on the log-transformed viral concentration (viral DNA copy number per total ng of DNA in the sample) of infected *Rana aurora* hosts. We refer to viral concentration as viral load for clarity. The median and 95% credible interval are shown for the coefficients in the linear model. Because there is no baseline effect of ATV included in these models due to zero ATV-infected individuals, the effects of co-exposure are compared to the single dose cases. Thus, the meaningful effect of the ATV:FV3 co-exposure shows that this treatment led to higher viral loads compared to the single-dose FV3 treatment. ^a^The effect of double dose on FV3 viral load was marginal, whereby the 94.4% CI does not overlap zero (i.e. an *α* = 0.056 in frequentist statistics).

| Effect |  | Median Coefficient | 95% CI |
| --- | --- | --- | --- |
| Baseline viral load (i.e. average across all cases) | Intercept | 0.680 | (-4.30, 5.76) |
| Effect of mortality | **Died** | **5.55** | **(3.89, 7.16)** |
| Effect of each virus, compared to baseline | FV3 | -2.18 | (-8.03, 2.43) |
|  | RFV3 | -2.78 | (-7.74, 3.40) |
| Effects of double dosage, compared to single dosage | *Double:FV3* | 3.23 | (-0.08, 6.59)<sup>a</sup> |
|  | **Double:RFV3** | **1.78** | **(0.01, 6.2)** |
| Effects of co-exposure, compared to single dosage | FV3:RFV3 | -0.706 | (-6.17, 4.8) |
|  | ATV:RFV3 | 0.166 | (-2.48, 2.96) |
|  | **ATV:FV3** | **3.39** | **(1.01, 5.78)** |

We successfully sequenced viral DNA from all but three of the infected individuals. The three individuals from which we did not successfully sequence had the three lowest viral loads. Sequencing results revealed that infected individuals were predominantly infected by the FV3-like strains. However, because FV3 and R-FV3 are indistinguishable based on this sequencing method, we could not reliably determine whether coinfection occurred (i.e. simultaneous presence of multiple virus types). Interestingly, four sequences from the R-FV3 single-virus exposures and the ATV + R-FV3 exposures showed 100% sequence identity to one another but did not perfectly match the sequences of the three viruses used in this experiment. We searched for similar sequences on GenBank^®^ via BLAST, which revealed a 100% match to an isolate of FV3 discovered in lungless salamanders of the Great Smokey National Park, TN (Gray et al. 2009a). The source of this contamination – whether two viruses were co-isolated from the bull frog culturing facility, or whether the original R-FV3 stock was contaminated post-culturing – is unclear, but it is unlikely to have affected our results.

### Experiment 2

No *Pseudacris triseriata* individuals died in the control group. Across all treatments, no individuals had died by 4 days post-exposure (dpe). Overall, only four individuals died in the ATV-only treatment (5.5% of all replicate individuals), and 12 (16%) and 14 (19%) individuals died in the FV3+ATV and FV3-only treatment replicates, respectively. When testing for an effect of co-exposure on survival rates, we found no overall difference between the survival rates in the FV3-only and FV3+ATV treatments (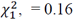, p = 0.68), although there was significant variation in survival rates among replicates (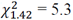, p = 0.038).

No individuals in the control treatment became infected; however, unlike *R. aurora*, which showed no infectivity with ATV, three *P. triseriata* individuals in the ATV-only treatment became infected, which were detected in the three different replicates (one per replicate) at 21dpe. Treatment and time post-exposure interacted to drive infection prevalence (Fig. 2; Table 3); for the FV3-only treatment, the proportion of infected individuals increased more consistently and substantially over time compared to the FV3+ATV treatment (Fig. 2). All three replicates of the FV3-only treatment at 21dpe had the same proportion of individuals infected (8/12, 66%). The FV3+ATV treatment replicates had more variable prevalence. In one of the FV3+ATV treatments at 4dpe, 9/12 (75%) of individuals were infected, which was a substantially larger proportion compared to all other 4dpe replicates, and the highest prevalence in the experiment overall (Fig. 2).

**Table 3.** Effects of time and co-exposure on the prevalence of infection in experimental *Pseudacris triseriata* populations. As in Table 1, the median and 95% credible interval are shown for the coefficients in the logistic model. The meaningful interaction was driven by a more substantial increase in prevalence over time in the FV3-only treatment group (Fig. 2).

| Effect |  | Median Coefficient | 95% CI |
| --- | --- | --- | --- |
| Time effects (factors) | 4dpe | -0.329 | (-10.26, 9.56) |
|  | 21dpe | 3.88 | (-6.24, 14.12) |
| Treatment effects | FV3 | 0.643 | (-9.25, 10.62) |
|  | FV3+ATV | 3.23 | (-7.18, 13.68) |
| Interaction effect | **Time x Treatment** | **-1.90** | **(-3.43, -0.45)** |

**Figure 2:**
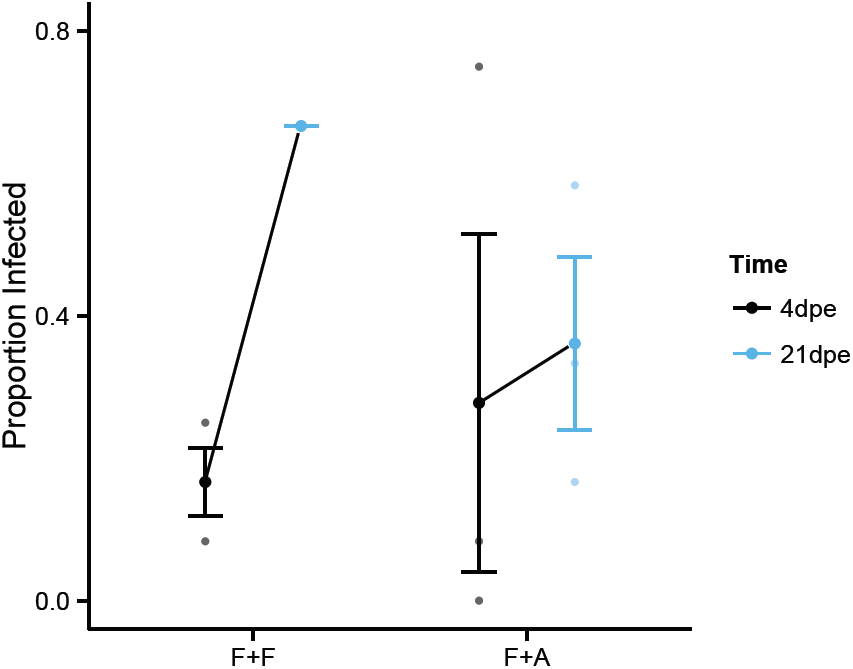
Proportion of *Pseudacris triseriata* infected individuals between time points and treatments in experiment 2. Time point are distinguished by color, as depicted in the legend. Large, bold circles represent the mean prevalence, and error bars represent one standard error of this mean. Smaller and more opaque circles represent the prevalence of the replicate larval populations. Note that all 3 replicates of the FV3-only treatment at 21dpe had the same prevalence.

Individuals that died in the experiment had, on average, higher viral loads compared to infected individuals that were sampled prior to mortality (Fig. 3; Table 4). Because all of the individuals that died were in the 21dpe treatments, the average viral load increased between the 4dpe and 21dpe treatments (Fig. 3). There were no overall effects of co-exposure on viral load in this experiment (Table 4).

**Table 4.** Effects of time and co-exposure on the prevalence of the viral concentration (viral load) in infected *Pseudacris triseriata*. The median and 95% credible interval are shown for the coefficients in the linear model. In this model we included a random effect of replicate population on the intercept, and therefore the random effect standard deviation and residual standard deviation are shown.

| Effect |  | Median Coefficient | 95% CI |
| --- | --- | --- | --- |
| Baseline viral load (i.e. average across all cases) | Intercept | 4.35 | (-3.61, 12.42) |
| Effect of mortality | **Died** | **3.01** | **(0.17, 5.82)** |
| Time effects (factor) | 4dpe | -1.69 | (-7.84, 4.5) |
|  | 21dpe | 0.716 | (-6.66, 8.3) |
| Treatment effects | FV3 | 1.33 | (-5.11, 7.71) |
|  | FV3+ATV | -2.16 | (-9.6, 5.77) |
| Interaction | Time x Treatment | 0.724 | (-2.92, 4.2) |
| Standard deviation among replicates | $\sigma_{\text{replicate}}$ | 1.40 | (0.15, 3.45) |
| Residual standard deviation | $\sigma_{\text{residual}}$ | 3.88 | (3.2, 4.86) |

**Figure 3:**
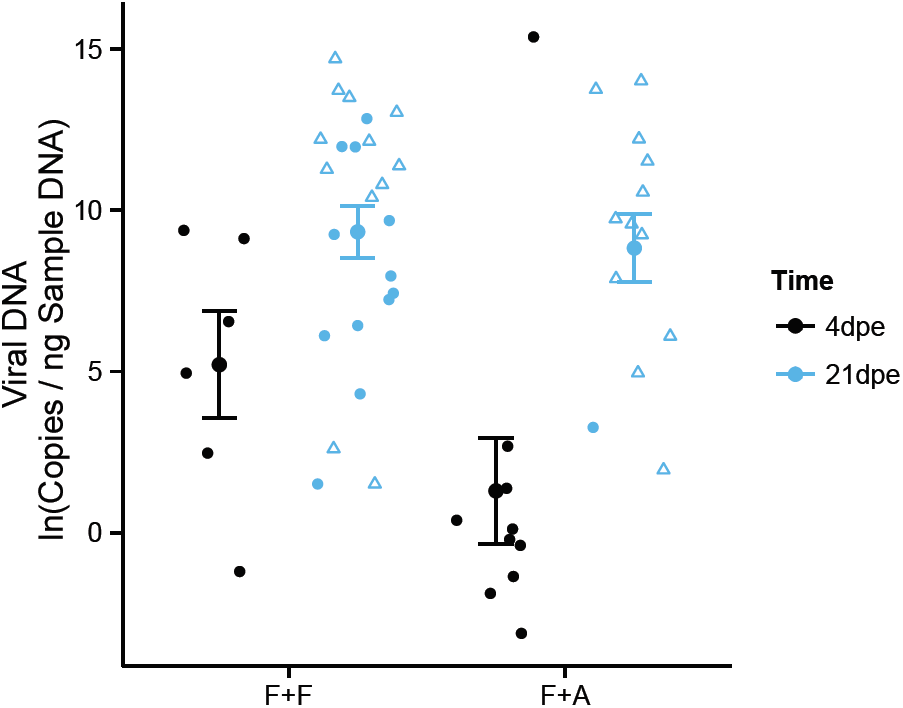
Viral copy number across time points and treatments in experiment 2 with *Pseudacris triseriata*. ATV-only treatments are not shown, because only 3 total individuals became infected, all in the 21dep treatments (one individual per replicate). Time points are distinguished by color, as depicted in the legend. Large, closed circles represent the mean viral load, and error bars represent one standard error of the mean. Smaller closed circles represent the viral load of infected individuals that survived until the end of the experiment (or until destructive sampling in the case of 4dpe replicates). Open triangles represent infected individuals that died prior to the end of the experiment. A jitter is added to the data for ease of interpretation. Notice the most heavily infected individual from the FV3+ATV treatment at 4dpe.

We amplified and sequenced viral DNA from 85% of the infected individuals (n = 45 / 53). Of the three individuals that tested positive for infection in the ATV-only treatments, two DNA samples amplified, and their sequence data matched that of ATV, verifying that *P. triseriata* can become infected with ATV. All sequences from the FV3-only treatment matched FV3 DNA. Only one individual from the FV3+ATV treatment was infected with ATV, which was also the host with the highest observed viral load (4.76 x 10^6^ viral DNA copies ng^−1^ DNA) and came from the replicate population with the highest infection prevalence (75% in the FV3+ATV, 4dpe treatment).

## Discussion

We conducted experiments to identify the effects of co-exposure to multiple ranaviruses at the scale of both individual hosts and experimental populations. For individual hosts, co-exposure to *Ambystoma tigrinum* virus (ATV) and frog virus 3 (FV3) increased the infection success of FV3. However, this same effect did not hold for co-exposure to two more related strains (FV3 and *Rana catesbeiana* virus (RCV-Z2), herein R-FV3), indicating that viral identity and viral relatedness may be important for predicting the outcome of co-exposure. At the host population-scale, we found some evidence that, when ATV co-occurs with FV3, co-exposure can lead to higher infection prevalence in the population. By conducting experiments at both the individual- and population-level scales, results of this study indicate that the co-occurrence of *Ranavirus* species has the potential to alter epizootic dynamics in natural amphibian populations.

In our first experiment, in which we exposed individual *Rana aurora* to multiple ranaviruses, we demonstrated several expected results that help validate our methods and experimental design. First, we found higher infection prevalence in the double-dose FV3 treatment, compared to the single-dose FV3 treatment, showing that our chosen differences in dose led to measurable differences in infectivity. We also found higher viral loads in individuals that died compared to surviving individuals. This result is intuitive, especially considering evidence that the virulence of ranaviruses is at least partially associated with within-host viral replication (Brunner & Collins 2009). We also show that ATV is not highly infectious in larval frogs, corroborating previous findings (Jancovich et al. 2001, Schock et al. 2008); however, we demonstrate that *Pseudacris triseriata* is susceptible to this virus in our second experiment. Although mortality was high in our first experiment, the mortality patterns were consistent across treatments, and we believe that the basic results outlined above show that our methods were unbiased.

Critically, our first experiment provides evidence for an effect of co-exposure on viral infectivity and viral replication. The data suggest that co-exposure to ATV and FV3 synergistically increased the host’s probability of infection with FV3. This effect of co-exposure with ATV was seen with two FV3-like strains, wild-type FV3 and R-FV3. Notably, we saw this co-exposure effect with ATV+R-FV3, even though we were unable to accurately quantify the titer of the R-FV3 stock, which demonstrates a robust effect of co-exposure on prevalence. We also saw that co-exposure to ATV and FV3 led to higher average viral load compared to the single-dose FV3 treatment. Given that no individuals became infected with ATV alone, and that the ATV+FV3 co-exposure constitutes a single dose of FV3, this latter result suggests a synergistic effect of co-exposure on viral replication within a host. We believe our results imply that ATV and FV3 likely either coinfected or superinfected the hosts and that exposure to ATV facilitated the invasion and subsequent proliferation of FV3 within larvae (discussed more below).

Our second experiment, in which we exposed experimental populations of *P. triseriata* to multiple ranaviruses, provides some additional evidence for the effect of co-exposure on epizootics. Although we detected ATV infection in only one individual in the ATV+FV3 treatment group, from one replicate population, this individual exhibited the highest overall viral load in our experiment, even after only 4dpe. This individual also was sampled from the replicate population with the largest epizootic in the experiment (75%). This evidence, in combination with the result that ATV only rarely infects these frogs, suggests that when ATV is able to establish infections in a population concurrent with FV3, there is the potential for larger epizootics.

We suspect that the effect of co-exposure was not as strong in the second experiment because of the difference in viral delivery and dosage, or perhaps host species identity. Specifically, ATV was only able to establish in one replicate co-exposure population (as evidenced by our sequencing methods), and we thus only saw one FV3+ATV co-exposure population with an effect. There are several plausible explanations for this outcome. First, the majority of pre-exposed individuals may not have become infected, and therefore the susceptible, replicate populations were never exposed to ATV. Second, the pre-exposed individuals were infected, but cleared the infection prior to the population phase of the experiment. Or third, the pre-exposed individuals were infected with ATV but cleared the infection during the population phase of the experiment before ATV could infect other susceptible individuals. Because we were not successful in marking pre-exposed individuals, we cannot distinguish between these scenarios.

In the first experiment, each co-exposed individual was passively exposed to ∼1×10^6^ PFU of ATV in solution. However, in the second experiment, the susceptible larvae in the replicate populations could only become exposed to ATV if the pre-exposed individuals were infectious. Thus, it is likely that if we had passively exposed the replicate populations to ATV in a way similar to our first experiment, a larger effect of co-exposure would be seen. The differences in effects between the two experiments could also be due to differences in the effects of co-exposure among amphibian species. It is known that variability in FV3 infectivity among amphibian species has phylogenetic and ecological correlates (Hoverman et al. 2010, 2011).

Somewhat surprisingly, we did not see an effect of co-exposure on overall survival rates in our experiments. We ran our experiments for 21d, which in previous studies has been long enough to see 20-100% mortality due to ranavirus infection in other species of frogs and salamanders (Brunner et al. 2005, Hoverman et al. 2010). Given that case-mortality rates tend to be high (> 90%) for ranaviruses, it is likely that more individuals, especially in the co-exposure treatments with higher viral loads, would have died due to infection if we carried out the experiments for a longer time period.

Based upon the evidence from the first experiment, we propose two hypotheses for the observed increase in infectivity and viral replication following host co-exposure to ATV and FV3. First, exposure to these two distinct virus types could lead to non-overlapping immune responses in the amphibian larvae, which leads to a trade-off that decreases the efficacy of the host’s response to FV3, facilitating invasion. While there is ample evidence for resource competition in multi-strain infections (Read & Taylor 2001, Mideo et al. 2008, Alizon et al. 2013b), few studies have documented the possible immune trade-offs imposed by multi-strain infections (Balmer & Tanner 2011). In the *Ranavirus* system, along with complex innate immune responses, *Xenopus laevis* adults produce long-lasting anti-FV3 IgY antibodies, and larvae produce less effective innate and adaptive responses (Chinchar et al. 2011, Chen & Robert 2011). However, it is unknown if exposure to ATV elicits overlapping innate and adaptive responses with FV3. Future experiments that determine the degree of antibody specificity between ATV and FV3 and that alter the timing of exposure between FV3 and ATV may help to further test this hypothesis of the effect of co-exposure.

A second, alternative hypothesis for the effect of co-exposure is viral recombination. It is possible that, if ATV and FV3 coinfect the same host cells, recombination could occur to produce a novel, more infectious virus. Genomic evidence from multiple *Ranavirus* species suggests high recombination frequency and shows that these viruses are prone to host-shifts due to gene acquisition and subsequent adaptation (Jancovich et al. 2003, 2010, Abrams et al. 2013). Recombination has been employed to explain the collinearity and the one inversion between the ATV and FV3 genomes (Eaton et al. 2007). Furthermore, it was recently discovered that the R-FV3 strain we used here (RCV-Z2) is the product of a recombination event between an FV3-like strain and a common midwife toad virus (CMTV)-like strain from Europe, and this recombination is likely the cause of the high virus-induced mortality rate of this strain (Claytor et al. 2017). This hypothesis of recombination could be tested by isolating many viruses from the co-exposure group via plaque assay, growing the viruses in culture, and conducting full genome sequencing and alignment to both FV3 and ATV.

Our results illustrate that in natural amphibian populations, co-occurrence of ATV and FV3 could alter epizootic dynamics. Specifically, if ATV can establish in a larval frog population, co-occurrence with FV3 could result in more infected individuals and subsequently higher mortality rates in the long run. This effect seems particularly relevant for wetlands in which salamanders and frogs cohabitate. If ATV is present and infects the local salamanders and FV3 establishes in the anuran populations, spillover of ATV from the salamanders could enhance FV3 epizootics in the frogs. Also, because FV3 is adept at infecting salamanders as well (Schock et al. 2008), it is likely that such a scenario would increase infection prevalence and intensity in the urodele population. Thus, our results illustrate the need to consider co-exposure and co-infection in the amphibian-*Ranavirus* system and emphasize the need for field data on ATV and FV3 co-occurrence at both the wetland- and host individual-levels.

This study adds to a growing body of literature that illustrates the important consequences of multi-pathogen interactions in mediating pathology and transmission. Furthermore, our results emphasize the importance of multi-scale experiments for understanding how interactions among pathogens influence transmission. In general, the impact of co-exposure on transmission will depend on how pathogen interactions within hosts feedback on between-host dynamics. In the *Ranavirus* system, co-exposure increased pathogen infection success and viral replication within hosts but did not result in more rapid host death, ultimately leading to increased transmission when both pathogens co-occurred. Research that integrates multi-scale experiments across a variety of systems will help us better understand the conditions under which co-exposure will significantly impact epidemics and epizootics.

## Acknowledgements

We would like to thank the Johnson Laboratory at CU-Boulder, particularly MB Joseph, for their helpful comments on a previous version of this manuscript. We are especially grateful to the hard work of two laboratory technicians, C. Ramsay and B. Camenga. This research was supported by the NSF Graduate Research Fellowship Program (DGE 1144083), the NSF (DEB-1149308) and NIH (R01GM109499), a Beverly Sears Graduate Student Research Grant, CU EBIO Departmental Research Grants, and the CU Natural History Museum Grant Program.

---

**Figure A1:**
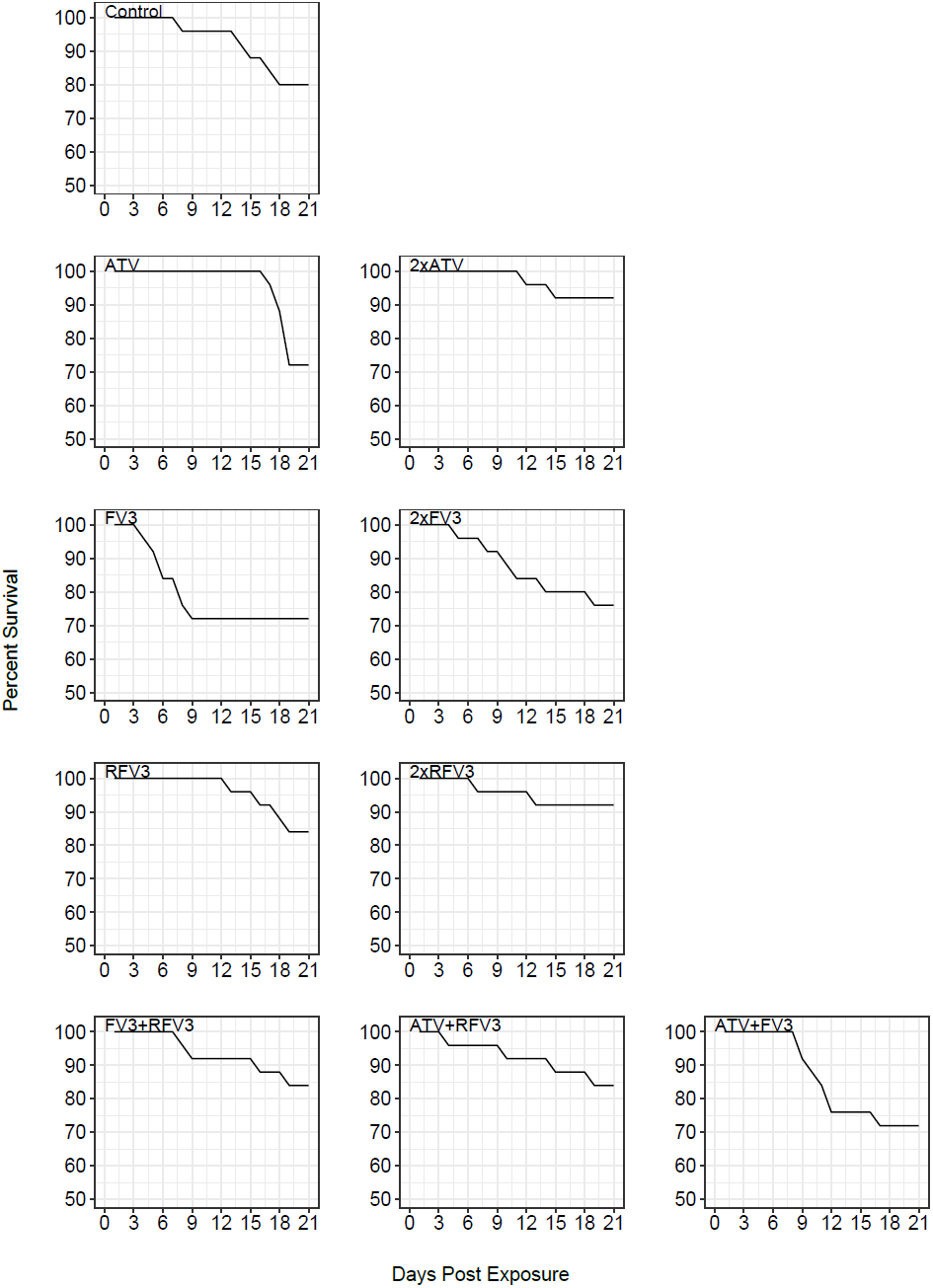
Survival curves for all treatments in experiment 1 with *Rana aurora.*

**Figure A2:**
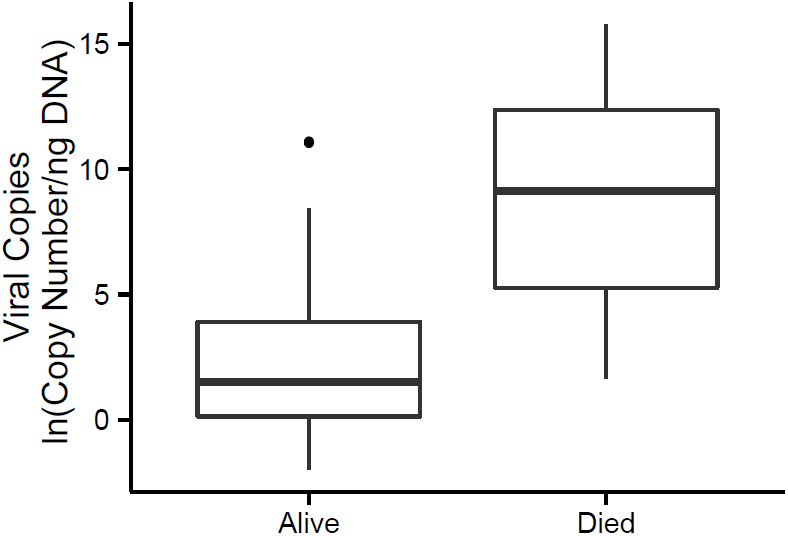
Boxplots (as in Figure 2) of viral load for infected *Rana aurora* that survived until the end of the experiment (Alive) or that died due to infection (Died).

